# Alpha variant versus D614G strain in the Syrian hamster model

**DOI:** 10.1101/2021.04.19.440435

**Authors:** Maxime Cochin, Léa Luciani, Franck Touret, Jean-Sélim Driouich, Paul-Rémi Petit, Grégory Moureau, Cécile Baronti, Caroline Laprie, Laurence Thirion, Piet Maes, Robbert Boudewijns, Johan Neyts, Xavier de Lamballerie, Antoine Nougairède

**Author notes:** These authors contributed equally to this article (Maxime Cochin, Léa Luciani). **Corresponding author:** Antoine Nougairède.

## Abstract

Late 2020, SARS-CoV-2 Alpha variant from lineage B.1.1.7 emerged in United Kingdom and gradually replaced the G614 strains initially involved in the global spread of the pandemic. In this study, we used a Syrian hamster model to compare a clinical strain of Alpha variant with an ancestral G614 strain. The Alpha variant succeeded to infect animals and to induce a pathology that mimics COVID-19. However, both strains replicated to almost the same level and induced a comparable disease and immune response. A slight fitness advantage was noted for the G614 strain during competition and transmission experiments. These data do not corroborate the epidemiological situation observed during the first half of 2021 in humans nor reports that showed a more rapid replication of Alpha variant in human reconstituted bronchial epithelium.

## Introduction

The genetic evolution of the severe acute respiratory syndrome coronavirus 2 (SARS-CoV-2) virus is a constant concern for medical and scientific communities. From January 2020, viruses carrying the spike D614G mutation emerged in several countries[1–3]. In June, D614G SARS-CoV-2 lineage B.1 became the dominant form of circulating virus worldwide and replaced the initial SARS-CoV-2 strains related to the outbreak in Wuhan, China. Experimental data from human lung epithelium and animal models revealed that the D614G substitution increased virus infectivity and transmissibility as compared to an original D614 strain[4]. However, it seems that this G614 variant does not cause more severe clinical disease. In late 2020, three SARS-CoV-2 variants sharing the N501Y spike mutation located in the receptor binding motif (RBM) emerged almost simultaneously in the United Kingdom (Alpha variant from lineage B.1.1.7; initially named VOC 202012/01)[5], in South Africa (Beta variant from lineage B.1.351)[6] and in Brazil (P.1 variant from lineage B.1.1.28.1)[7]. As previously observed with the G614 strain, the Alpha variant spread rapidly and became dominant in United Kingdom in December 2020, and in many other European and non-European countries from February 2021 onwards[8]. The Alpha variant harbors 8 additional spike mutations, including substitutions and deletions, compared to G614 circulating strains. From May 2021, a new variant that appeared in India, the Delta variant (lineage B.1.167.2), spread suddenly throughout the world, totally surpassing Alpha variant. This Delta variant possesses 8 spike mutations in comparison with G614 strains. The last variant of concern recognised by the WHO is the Omicron (lineage B.1.1.529) variant which appeared in November 2021 in Africa and which bears 32 spike mutations in comparison with G614 strain. The Omicron variant is currently spreading in many countries and is strongly suspected to become the new dominant variant. The regular emergence of variants that become world dominant in a few months, overtaking previous variants, seems to be associated with an improved affinity of the viral spike protein for the human angiotensin-converting enzyme 2 (ACE2) receptor[9,10]. In addition, the public health strategy to control the COVID-19 pandemic is currently based on the massive distribution of vaccines throughout the world. These efforts have been successful in reducing the number of infections and the burden of COVID-19 waves. However, all currently available approved vaccines were developed from the genetic sequence of the spike protein of the original virus that emerged in Wuhan in 2019. The issue now is whether the regular emergence of new variants with multiple mutations in the spike protein will compromise the effectiveness of the vaccine strategy.

We recently described the fitness advantage of Alpha variant using a model of reconstituted bronchial human epithelium[11]. In the present work, we compared the phenotype of the Alpha variant (hCoV-103 19/Belgium/rega-12211513/2020 strain) with that of a G614 strain (Germany/BavPat1/2020 strain) in the Syrian hamster (*Mesocricetus auratus*) model. The study includes comparison of viral replication kinetics, transmissibility, lung pathology, clinical course of the disease and immunological response.

## Materials and methods

### Cells and viruses

VeroE6 cells (ATCC CRL-1586) were grown at 37°C with 5% CO_2_ in minimal essential medium (MEM) supplemented with 1% non-essential amino acids, 1% Penicillin/Streptomycin and 7% of heat-inactivated fetal bovine serum (FBS) (all from ThermoFisher Scientific). VeroE6/TMPRSS2 cells (NIBSC 100978) were grown in the same medium supplemented with 2% of G-418 (ThermoFisher Scientific).

All experiments with infectious virus were conducted in biosafety level (BSL) 3 laboratory.

The SARS-CoV-2 strain BavPat1/2020 (G614 strain), supplied through European Virus Archive GLOBAL (European Virus Archive Global # 026 V-03883), was kindly provided by Christian Drosten (Berlin, Germany). The SARS-CoV-2 Alpha variant (lineage B.1.1.7) hCoV104 19/Belgium/rega-12211513/2020 strain (EPI_ISL_791333), used for *in vivo* experiments, was isolated from a naso-pharyngeal swab from a traveler returning to Belgium in December 2020. The SARS-CoV-2 BetaCoV/France/IDF0372/2020 strain (D614 strain) was supplied through European Virus Archive Global (European Virus Archive Global # 014V-03890). Virus stocks of these strains were produced using VeroE6 cells (passage history: 2 for G614 strain and Alpha variant, 3 for D614 strain). The SARS-CoV-2 Alpha variant (lineage B.1.1.7), hCoV-19/France/PAC-7b-exUK/2021 strain (EPI_ISL_918165), used for seroneutralization assays, was isolated from a 18 years-old patient. This strain is available through European Virus Archive Global (European Virus Archive Global # 001V-04044). The SARS-CoV-2 Beta variant (lineage B.1.351), hCoV-19/France/PAC-1299/2021 strain (EPI_ISL_1834082), was isolated from a naso-pharyngeal swab in France in 2021. This strain is available through European Virus Archive Global (European Virus Archive Global # 001V-04067). The SARS-CoV-2 Delta variant (B.1.617.2), hCoV-19/France/PAC-0610/2021 strain (EPI_ISL_2838050), was isolated from a 87 years-old patient in France in 2021. This strain is available through European Virus Archive Global (European Virus Archive Global # 001V-04282). Virus stocks of these variant were produced using VeroE6 TMPRSS2 cells (passage history: 2 for Alpha variant, 1 for Beta and Delta variants).

All virus stocks were characterized by full-genome sequencing (Ion Torrent) in order to verify the absence of additional mutations, especially in the spike-coding region when compared to sequences of seeded viruses.

### *In vivo* experiments

Following approval by the local ethical committee (C2EA—14) and the French ‘Ministère de l’Enseignement Supérieur, de la Recherche et de l’Innovation’ (APAFIS#23975), *in vivo* experiments were performed in accordance with the French national guidelines and the European legislation covering the use of animals for scientific purposes.

For each experiment, groups of three-week-old female Syrian hamsters (Janvier Labs) were intranasally infected under general anesthesia (isofluorane) with 50μL containing 2×10^3^, 10^4^ or 20 TCID_50_ of virus diluted in 0.9% sodium chloride solution. Mock-infected animals were intranasally inoculated with 50μl of 0.9% sodium chloride solution. Comparative and competition experiments were performed twice (two independent experiments with groups of 6 animals). Pooled data from both experiments (12 animals) were presented. Groups of 4 animals were used for the histology experiment. Groups of 6 to 12 animals were used for transmission experiments. Animals were maintained in ISOcage P - Bioexclusion System (Techniplast) with unlimited access to water/food and 14h/10h light/dark cycle. Animals were monitored and weighed daily throughout the duration of the study to detect the appearance of any clinical signs of illness/suffering. Nasal washes were performed under general anesthesia (isoflurane). Blood and organs were collected after euthanasia (cervical dislocation; realized under general anesthesia (isofluorane)).

### Organ collection

Nasal washes were performed with 150μl 0.9% sodium chloride solution which was transferred into 1.5mL tubes containing 0.5mL of 0.9% sodium chloride solution, then centrifuged at 16,200g for 10 minutes and stored at −80°C. Lung, gut and blood samples were collected immediately after the time of sacrifice. Left pulmonary lobes were washed in 10mL of 0.9% sodium chloride solution, blotted with filter paper, weighed and then transferred into 2mL tubes containing 1mL of 0.9% sodium chloride solution and 3mm glass beads. Guts (part of small and large bowels) were empty of their alimentary bolus, weighed and then transferred into 2mL tubes containing 1mL of 0.9% sodium chloride solution and 3mm glass beads. Left pulmonary lobes and guts were crushed using a Tissue Lyser machine (Retsch MM400) for 20min at 30 cycles/s and then centrifuged 10min at 16,200g. Supernatant media were transferred into 1.5mL tubes, centrifuged 10 min at 16,200g and stored at −80°C. One milliliter of blood was harvested in a 2mL tube containing 100μL of 0.5M EDTA (ThermoFischer Scientific) and then centrifuged 10 min at 16,200g to obtain plasma. Serum samples were collected from blood harvested in a 2mL tube incubate 15 min at room temperature and then centrifuged 10 min at 16,200g. Blood-derived samples were stored at −80°C. To assess the expression level of seven cytokines in lungs, right apical lobes were collected into 2mL tubes containing 0.75mL of Qiazol lysis reagent (Qiagen) and 3mm glass beads. They were then crushed using a Tissue Lyser machine (Retsch MM400) for 10min at 30 cycles/s and stored at −80°C.

### TCID_50_ assay

Virus titration was performed using 96-well culture plates containing confluent cells (VeroE6 cells, except for competition experiments between G614 and D614 strains where VeroE6 TMPRSS2 cells were used) inoculated with 150μL per well of four-fold dilutions of samples (dilution with medium supplemented with 2.5% FBS). After 6 days of incubation (37°C, 5% CO_2_) the absence or presence of cytopathic effect in each well was read and infectious titers were estimated using the Reed & Muench method[12].

### Molecular biology

For viral quantification, nucleic acids from each sample were extracted using QIAamp 96 DNA kit and Qiacube HT robot (both from Qiagen). Viral RNA yields were measured using a RT-qPCR assay targeting the *rdrp* gene as previously described[13].

To analyze samples from competition and transmission experiments (ie. with animal infected with a mix of both viruses), we used two specific RT-qPCR assays targeting the NSP6 coding region (each specifically detecting one of the competing viruses) to determine the proportion of each viral genome. Prior to PCR amplification, RNA extraction was performed as described above. RT-qPCR were performed with SuperScript III Platinum One-Step qRT-PCR kit (SuperScript™ III Platinum™ One-Step qRT-PCR kit, universal Invitrogen) using 2.5μL of nucleic acid extract and 7.5μL of RT-qPCR reagent mix. Using standard fast cycling parameters, i.e., 50°C for 15min, 95°C for 5min, and 40 amplification cycles (15 sec at 95°C followed by 45 sec at 55°C). RT-qPCR reactions were performed on QuantStudio 12K Flex Real-Time PCR System (Applied Biosystems) and analyzed on QuantStudio 12K Flex Applied Biosystems software v1.2.3. Viral RNA quantities were calculated using serial dilutions of T7-generated synthetic RNA standards. Primers and probes sequences were: Fwd: 5’-CATGGTTGGATATGGTTG-3’; Rev: 5’-GATGCATACATAACACAG-3’; Probe that specifically detect the G614 virus: 5’-FAM-GTCTGGTTTTAA-BHQ1-3’; Probe that specifically detect the 201/501YV.1 variant: 5’-VIC-TAGTTTGAAGCT-BHQ1-3’.

To quantify D614:G614 ratios using a previously described method[4], 498bp fragment that contained the spike mutation D614G was amplified from extracted RNA (QIAamp 96 DNA kit and Qiacube HT robot). RT-qPCR were performed with SuperScript III Platinum One-Step qRT-PCR kit (SuperScript™ III Platinum™ One-Step qRT-PCR kit, universal Invitrogen) using 5μL of nucleic acid extract and 20μL of RT-qPCR reagent mix using cycling parameters, i.e., 45°C for 30min, 94°C for 2min, and 40 amplification cycles (30sec at 94°C followed by 45 sec at 56°C and 2min at 72°C) followed by a last step at 72°C for 10min. RT-qPCR reactions were performed on 2720 Thermal cycler (Applied Biosystems). Primer sequences used for this first amplification were: Fwd: 5’-TGCACCAGCAACTGTTTGTGGACCT-3’and Rev: 5’-ACGTGCCCGCCGAGGAGAA-3’. The amplicons were then purified using NucleoFast 96 PCR Plate (Macherey-Nagel) coupled to a vacuum pump. Sequencing reactions using purified RT-PCR products were performed with BigDye Terminator v1.1 cycle sequencing kit (Applied Biosystems) using standard cycling parameters, i.e., 96°C for 1min and 25 cycles (10sec at 96°C, 5sec at 50°C and 3min at 60°C) and a 2720 Thermal cycler (Applied Biosystems). For each RT-PCR product, two sequencing reactions were performed using the following primers: Fwd: 5’-GGTTTAACAGGCACAGGTGTTCTTACTGAG-3’ and Rev: 5’-CTAGCGCATATACCTGCACCAATGGG-3’. The sequencing reactions were purified using Sephadex G-50 Medium (Cytivia) and analyzed on a 3500XL Genetic Analyser (Applied Biosystems). The proportion of electropherogram peak height representing mutation site of each competition was analysed using QSVanalyser program[14]. To calculate the amounts of each virus present in our samples, we calculated the average proportion of peak heights obtained for each nucleotide with the two primers. We then multiplied the RNA copy number given by the RDRP-based real-time quantitative PCR by this average proportion. Consistency and accuracy of this competition assay were validated using RNA extracts from D614 and G614 viruses mixed at ratios of 10:0, 9:1, 7:3, 5:5, 3:7, 1:9 and 0:10 (Supplemental Table 1).

To assess the expression level of seven cytokines in lungs, 150μL of chloroform was added to crush lung samples. After, centrifugation (15min at 4°C and 12000g), 300μL of the aqueous phase was used for RNA extraction using EZ1 RNA tissue mini kit and EZ1 advanced XL robot (both from Qiagen). Quantitative RT-qPCR were performed using primers previously described by Dias de Melo *et al* [15] using QuantiNova SYBR^®^ Green RT-PCR Kit (Qiagen) according to the manufacturer instructions. RT-qPCR reactions were performed on QuantStudio 12K Flex Real-Time PCR System (Applied Biosystems) and analyzed on QuantStudio 12K Flex Applied Biosystems software v1.2.3. All cytokine mRNA quantifications were performed using serial dilutions of synthetic standards. Data were normalized to γ-actin reference gene relative expression[15].

### Seroneutralization assay

One day prior to infection, 5×10^4^ VeroE6/TMPRSS2 cells per well were seeded in 100μL assay medium (containing 2.5% FBS) in 96 well culture plates. The next day, 25μL of a virus mix diluted in medium was added to the wells. The amount of virus working stock used was calibrated prior to the assay, based on a replication kinetics, so that the viral replication was still in the exponential growth phase for the readout as previously described[16,17]. This corresponds here to a MOI of 0.002. Then six 2-fold serial dilutions of hamster sera starting at 1/10 were added to the cells (25μL/well, in assay medium) in duplicate. In addition, three 2-fold dilutions of a negative serum (from uninfected animals) starting at 1/10 were added to the plate to assess virus replication in presence of hamster serum (called ‘negative serum’ below). Four ‘virus control’ wells were supplemented with 25μL of assay medium to verify viral replication without serum. Plates were first incubated 15min at room temperature and then 2 days at 37°C prior to quantification of the viral genome by real-time RT-qPCR as previously described[18]. Briefly, nucleic acid from 100μL of cell supernatant were extracted using QIAamp 96 DNA kit and Qiacube HT robot (both from Qiagen). Viral RNA was quantified by real-time RT-qPCR (GoTaq 1 step RT-qPCR kit, Promega). Quantification was provided by serial dilutions of an appropriate T7-generated synthetic RNA standard. RT-qPCR reactions were performed on QuantStudio 12K Flex Real-Time PCR System (Applied Biosystems) and analyzed using QuantStudio 12K Flex Applied Biosystems software v1.2.3. Primers and probe sequences, which target SARS-CoV-2 N gene, were: Fw: 5’-GGCCGCAAATTGCACAAT-3’; Rev: 5’-CCAATGCGCGACATTCC-3’; Probe: 5’-FAM-CCCCCAGCGCTTCAGCGTTCT-BHQ1-3’. Percentage of viral inhibition was calculated as follows: 100* (mean quantity for negative serum-sample quantity)/mean quantity for negative serum. The 90% and 99% inhibition dilution factor were determined using logarithmic interpolation as previously described[16,18].

### Histology

Animal handling and hamster infections were performed as described above in the “in vivo experiments” section. Left pulmonary lobes were harvested after intratracheal instillation of a 4% (w/v) formaldehyde solution and fixed 72h at room temperature with a 4% (w/v) formaldehyde solution and then embedded in paraffin. Histological analysis was performed as previously described[13]. Briefly, 3.5-μm tissue sections were stained with hematoxylin-eosin (H&E) and analyzed blindly by a certified veterinary pathologist. Different lung compartments were examined: (1) bronchial and alveolar walls: a score of 0 to 4 was assigned based on the severity of inflammation; (2) alveoli: a score of 0 to 2 was assigned based on the presence and severity of hemorrhagic necrosis; and (3) vessel changes (leukocyte accumulation in the subendothelial space and tunica media): the absence or presence was scored 0 or 1, respectively. A cumulative score was then calculated and assigned to a severity grade (see Supplemental Table 2).

### Graphical representations and statistical analysis

Timelines (Figure 1.a and 2.a,b) were created on *biorender.com*. Graphical representations and statistical analysis were performed using GraphPad Prism 7 software (GrapPad software). For each group of data we applied the Shapiro-Wilk normality test. Then a two-by-two comparison of groups was performed using either an unpaired t test with or without a Welch’s correction if the variance did not assume an equal distribution (according to a Fisher test) or a Mann-Whitney test if the distribution was non-Gaussian. All statistical tests performed were two-sided when relevant.

**Figure 1:**
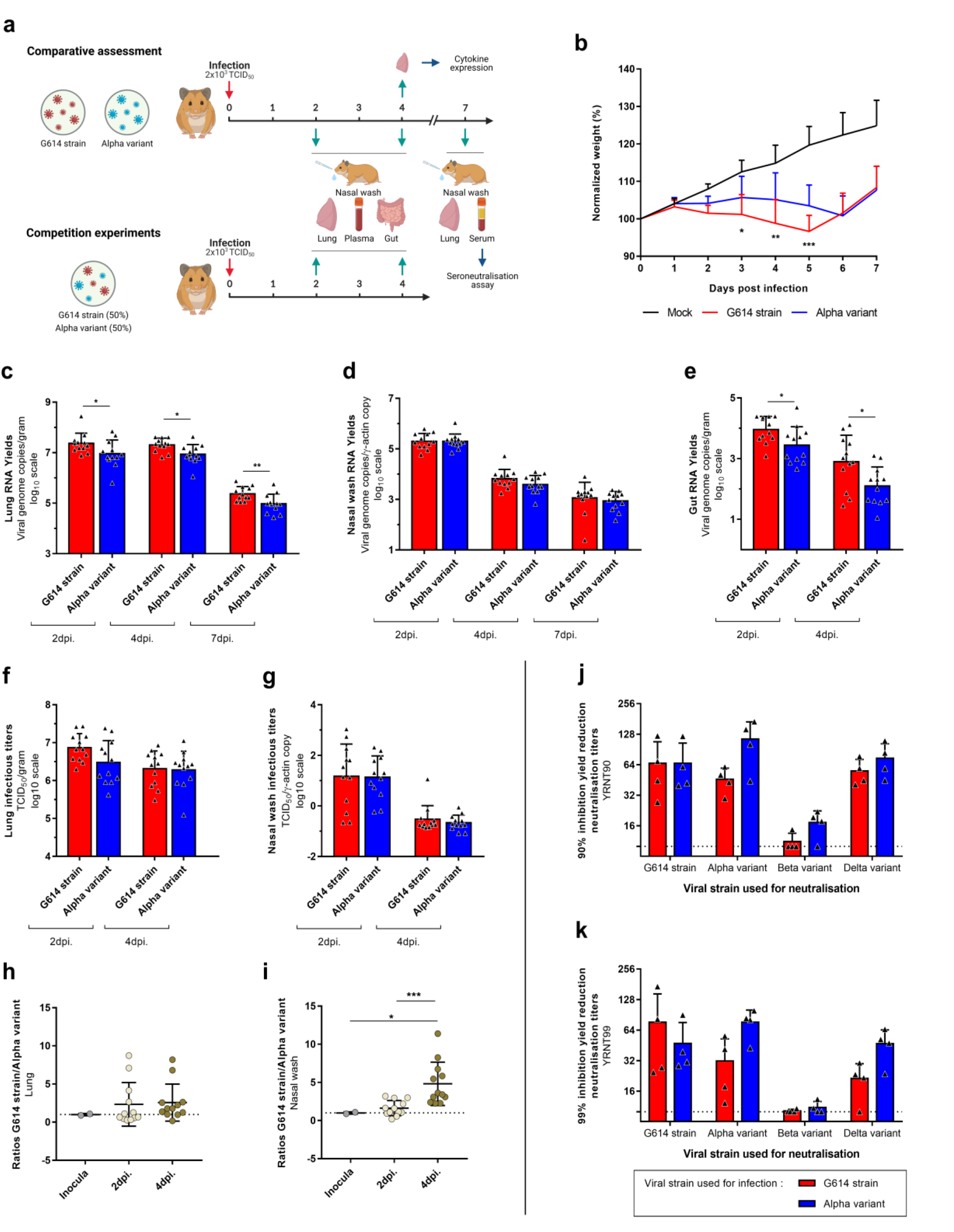
Clinical follow-up, viral replication in Syrian hamsters and seroneutralization tests. (a) Experimental timeline. Groups of 12 hamsters were intranasally infected with 2×10^3^ TCID_50_ of Alpha variant or G614 strain for comparative assessment, or with a mix (1:1) of both viral strains for competition experiment (10^3^ TCID_50_ of each). (b) Comparative clinical follow-up. Weights are expressed as normalized weights (i.e. % of initial weight). ***, ** and * symbols indicate that normalized weights for the Alpha variant group are significantly higher than those of the G614 group with a p-value ranging between 0.0001-0.001, 0.001–0.01, and 0.01–0.05, respectively (Two-way ANOVA test with Tukey’s post-hoc analysis). (c-e) Comparative assessment of viral RNA yields in lungs (c), nasal washes (d) and guts (e), measured using a RT-qPCR assay. ** and * symbols indicate that viral RNA yields for the Alpha variant group are significantly lower than those of the G614 group with a p-value ranging between 0.001–0.01, and 0.01–0.05, respectively (Mann-Whitney and Unpaired t tests). (f-g) Comparative assessment of infectious titers in lungs (f) and nasal washes (g), measured using a TCID_50_ assay. (h-i) Competition experiments. Two specific RT-qPCR assays were used to measure the quantity of each virus in lungs (h) and nasal washes (i). Results are expressed as [G614/ Alpha variant] ratios. *** and * symbols indicate that ratios at 4 dpi are higher than those at 2 dpi or in inocula with a p-value ranging between 0.0001-0.001 and 0.01–0.05, respectively (Mann-Whitney tests). (j-k) Seroneutralization tests performed with sera from animals sacrificed at 7 dpi. 90% (j) and 99% (k) Yield Reduction Neutralization Titers (90-99YRNT) were determined against four strains of SAR-CoV-2: G614 strain, Alpha variant, Beta variant and Delta variant. Results from statistical analysis are presented in Supplemental Table 4. (b-k) Data represent mean ± SD.

**Figure 2:**
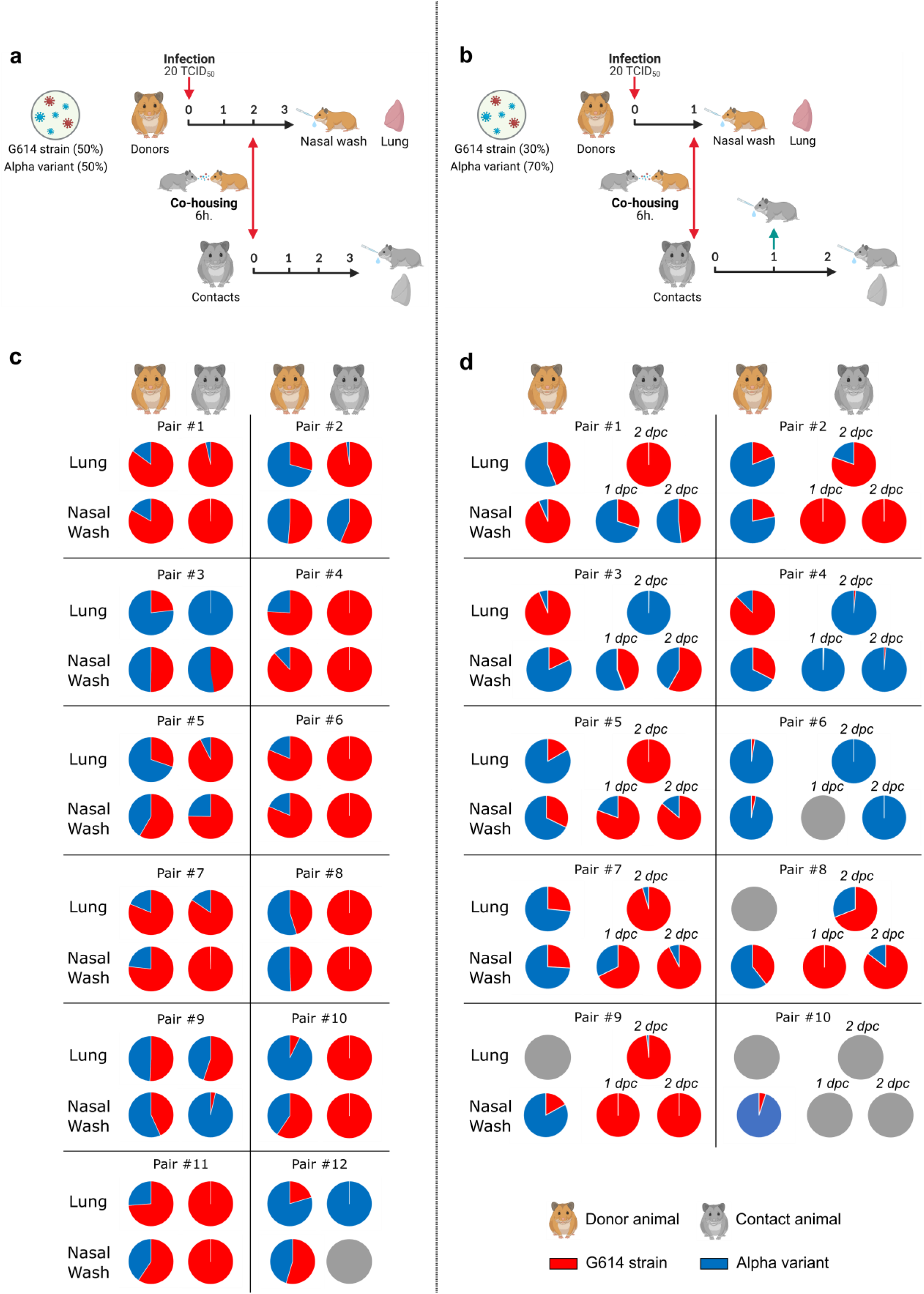
Transmissibility assessment. (a-b) Experimental timelines. (a) A group of 12 hamsters, named ‘donors’, was intranasally infected with an equal proportion of each viral strain for competition experiment (total: 20 TCID_50_). At 2 dpi, each donor was co-housed with a contact animal during a period of 6 hours. Donors and contacts were sacrificed at 3 dpi and at 3 dpc respectively. (b) A group of 10 hamsters, named donors, was intranasally infected with a mix (6:14) of G614 strain and Alpha variant for competition experiment (total: 20 TCID_50_). At 1 dpi, each donor was co-housed with a contact animal during a period of 6 hours. Donors and contacts were sacrificed at 1 dpi and at 2 dpc respectively. (c-d) Graphical representation of the proportion of each virus found in lungs and nasal washes for each pair of animals in transmission experimentations with 1:1 (c) and 6:14 ratios (d). Two specific RT-qPCR assays were used to measure the quantity of each virus. Grey circles mean that no viral RNA was detected in these nasal washes and lungs.

## Results

First, to detect modifications of the clinical course of the disease following infection with the Alpha variant, groups of 12 three-week-old female Syrian hamsters were intranasally infected with 50μl containing 2×10^3^ TCID_50_ of Alpha variant or G614 strain (Figure 1.a). Follow-up of these animals until 7 days post-infection (dpi) showed with both strains a lack of weight gain compared to mock-infected group. Normalized weights (i.e. % of initial weights) of animals infected with Alpha variant were significantly higher than those of G614 group at 3, 4 and 5 dpi (*p-values* between 0.0403 and 0.0007) (Figure 1.b). However, this difference seems to be the result of a delayed onset of disease for Alpha variant since significant difference of normalized weights compared to mock-infected group began at 2 dpi for animals infected with G614 strain and at 3 dpi for those infected with the Alpha variant.

Second, the lung pathological impairments induced by both strains was assessed in groups of four hamsters infected with 10^4^ TCID_50_ of virus (Supplemental Figure 1). Lungs collected at 5 dpi showed that both strains induced marked to severe pulmonary pathological changes without significant difference regarding cumulative scores (Supplemental Figure 1.a).

To further investigate viral replication, groups of 12 three-week-old female Syrian hamsters were intranasally infected with 50μl containing 2×10^3^ TCID_50_ of Alpha variant or G614 strain and several tissues were collected at different time points (Figure 1.a). Viral RNA quantification was performed using a RT-qPCR assay (i) in lung and nasal wash samples collected at 2, 4 and 7 dpi, and (ii) in blood and gut samples collected at 2 and 4 dpi. Infectious titers were determined using a TCID_50_ assay in lungs and nasal washes at 2 and 4 dpi. Overall, the results indicated that the Alpha variant properly replicate in the hamster gut and respiratory tract. However, higher viral RNA yields were found in all samples from animals infected with G614 strain (difference ranged from 0.085 to 0.801 log10). This difference was significant in lung and gut at any time point (*p values* ranging between 0.0332 and 0.0084) (Figure 1.c.d.e). Results of plasma did not show any significant difference (Supplemental Figure 2.a). A similar pattern was observed when assessing infectious viral loads using a TCID_50_ assay (differences ranged from 0.0343 to 0.389 log10) but no significant difference was found (Figure 1.f.g).

To detect more subtle differences of viral replication *in vivo*, we performed competitions experiments as previously described[9,19]. Groups of 12 animals were simultaneously infected intranasally with 50μl containing 50% (10^3^ TCID_50_) of each viral strain. Lungs, nasal washes and plasma were collected at 2 and 4 dpi (Figure 1.a). Using two specific RT-qPCR systems, we estimated in all samples the proportion of each viral genome in the viral population (expressed as G614 strain /Alpha variant ratios in Figure 1.h.i). Once again, results revealed that G614 strain seems to replicate a somewhat more efficiently and supplants progressively the Alpha variant. Indeed, G614 strain/Alpha variant estimated ratios at 4dpi were significantly higher than those at 2 dpi in nasal washes (*p*=0.0001). Moreover, ratios at 4 dpi in nasal washes were also significantly higher than those in the infecting inoculum (*p*=0.022) (Figure 1.i). By contrast, no significant difference was found in lungs (Figure 1.h) and plasma (Supplemental Figure 2.b).

To obtain a clearer picture, we compared the transmissibility of both strains in two different *in vivo* experiments.

During the first experiment, groups of 12 animals were simultaneously infected intranasally with 50μl containing a low dose of each viral strain (total: 20 TCID_50_). These animals, called ‘donors’, firstly housed individually, were co-housed at 2 dpi with an uninfected animal, called ‘contact’, during a period of 6 hours in a new cage. Then, donors returned in their initial cages and were sacrificed at 3 dpi. Contact animals were sacrificed at 3 days post-contact (dpc)(Figure 2.a). Using the two specific RT-qPCR systems used for competition experiments, we estimated in all samples (nasal washes and lungs) the proportion of each viral genome in the viral population (Figure 2.c). Data from lungs of donors showed for two animals, an equivalent proportion of both viruses (from 40% to 60% of each strain); for five animals, a majority (>60%) of G614 virus; and for the five remaining animals, a majority (>60%) of Alpha variant virus. However, we did not find the same distribution in nasal washes in which we observed: for eight animals, an equivalent proportion of both viruses; for four animals, a majority of G614 virus; and for no animal, a majority of Alpha variant virus. Consistently with this observation, we found a large majority (>75%) of G614 virus in lungs and nasal washes of eight contacts, and only two and one animals exhibited a large majority (>75%) of Alpha variant virus in lungs and nasal washes respectively (Figure 2.b). When analyzing the data from each pair of animals, we observed an increase of the proportion of G614 virus between the nasal wash of the donor and lungs of the contact in almost all cases (10/12). To confirm the suitability of this low-dose competition procedure, we applied it to compare the transmissibility of G614 and D614 strains (Supplemental Figure 3) since several studies already demonstrated that G614 strains were more transmittable than D614 strains[4,20]. We used for these experiments groups of 6 animals and a previously described protocol to estimate the proportion of each viral genome in the viral population[4]. As expected, we found a large majority of G614 strain in almost all lungs and nasal washes of contact animals at 3 dpc confirming the effectiveness of this method to compare the transmission of SARS-CoV-2 strains.

To offset the impact of the superior G614 replication on transmissibility assessment, we repeated a similar experiment in which we modified two parameters to co-house contact animals with donors carrying an equivalent proportion of both viruses in nasal washes as well as possible: donors were infected intranasally using a G614 strain:Alpha variant ratio of 6:14 (ie 30% and 70% of G614 strain and Alpha variant respectively) of and were co-housed with contact animals at 1 dpi. Moreover, we used group of ten animals and contact animals were sacrificed at 2 dpc in order to determine the dominant strain at early stage of infection. Nasal washes collected from donors right after the co-housing at 1 dpi showed for half of the animals a proportion of G614 strain ranging from 20 to 80%, for four animals a proportion of G614 strain ranging from 0 to 20% and for one animal a proportion of G614 strain upper than 80%. Of note, one contact (pair #10) was not infected during co-housing. Among the 8 contact animals co-housed with donors that carried a majority (>50%) of Alpha variant in nasal washes, 5 carried a large majority (>75%) of G614 strain in almost all samples (nasal washes at 1 and 2 dpc, and lungs at 2 dpc) while only 3 carried a large majority of Alpha variant. For the remaining contact animal co-housed with a donor that carried a majority of G614 strain (pair #1), both strains were found in nasal washes at equivalent level while only G614 strain was in majority in lungs.

Altogether, our results suggest that the replication of both G614 strain and Alpha variant were highly comparable in hamsters. Nonetheless, using a more sensitive method, we observed that the Alpha variant is outcompeted by the G614 strain; it results in an advantage for the G614 strain during transmission experiments. Notably, such results are not in line with experimental data *ex vivo* (human epithelial cultures grown at the air liquid interface) and with epidemiological observations.

We then compared transcriptional early immune signatures in lungs from animals sacrificed at 4 dpi following intranasal infection with 50μl containing 2×10^3^ TCID_50_ of Alpha variant or G614 strain. The expression level of seven cytokines (Interferon-γ, TNF-α, IL-6, IL-10, IL-1β, Cxcl-10, Ccl5) was quantified using RT-qPCR assays (Supplemental Figure 4; expressed as mRNA copies/γ-actin copies). Infection by both viral strains induced an important increase of CXCL10, CCL5, IFNγ, Il-6 and Il-10 expression levels (*p*<0.0001) and a moderate increase of Il-1β (*p*=0.0014 for G614 strain and *p*=0.0281 for Alpha variant) and TNFα (*p*=0.0389 for G614 strain and *p*=0.0350 for Alpha variant) expressions levels to mock-infected animals (Supplemental Figure 2). Comparison between animals infected with Alpha variant and G614 strain did not show any significant differences of cytokines expression levels. This suggests that the early immune response induced by both viral strains is similar, in line with a recent study that did not present major differences except an upregulation of Il-6, Il-10 and IFNγ with animals infected by Alpha variant[21].

Finally, we used sera collected at 7 dpi following intranasal infection with 50μl containing 2×10^3^ TCID_50_ of Alpha variant (n=4) or G614 strain (n=4)(Figure 1.a) to assess the level of protection against four circulating strains of SARS-CoV-2: the G614 strain, the Alpha variant, a ‘South-African’ Beta variant (lineage B 1.351) and an ‘Indian’ Delta variant (lineage 1.617.2). Sera were tested for the presence of antibodies using a 90-99% viral RNA Yield Reduction Neutralization Test (YRNT90/YRNT99) (Figure 1.j.k). Overall, results showed that animals infected with Alpha variant or G614 strains produced similar levels of neutralization antibodies against these strains with a slight advantage for animals infected with Alpha variant for YRNT99 titers against Alpha variant (*p* =0.0299) and Delta variant (*p* =0.0369). However, all infected animals produced lower neutralization titers against the Beta variant. This difference is significant with all animals when considering YRNT90 titers (*p values* between 0.0019 and 0.0450), and significant only with animals infected with Alpha variant when considering YRNT99 titers (*p*<0.0474) (Supplemental Table 4). This suggests an effective cross-immunity between Alpha variant, G614 strain and Delta variant but a reduced cross-protection against the Beta variant. These data indicate that only an active circulation of Beta variant might increase the risk of reinfection or failure of vaccination campaigns. This is in accordance with recently reported epidemiological observation[22–25].

## Discussion

Our results show that the Alpha variant induces a pathology that mimics human SARS-CoV-2 infection in the hamster model and can be used for preclinical analysis of vaccines and therapeutic agents. These data corroborate those of a recent study in which the same strains, a similar hamster model but higher virus inocula (10^5^ TCID_50_) were used[21]. Since its emergence in late 2020 in Europe, the Alpha variant spread across several continents and became the major circulating strain in many countries. Moreover, data from reconstituted human airway epithelia also showed a strong replicative fitness advantage for Alpha variant[11]. Notably, our findings in the hamster model are not in line with these observations. Of note *ex-vivo* models such as reconstituted human airway epithelia are less complex than *in vivo* model, especially in term of immune response and viral replication dynamic. Moreover, comparing fitness of strains that evolve in humans using a different species may be a significant bias factor. Indeed, recent studies suggest that the affinity of the SARS-CoV-2 spike protein for the ACE2 receptor is a species-dependent parameter[26,27]. Other unknown species-dependent mutations located in other genomic regions could also explain the observed discrepancy.

Some recent studies regarding transmissibility of Alpha variant or Alpha-like viruses showed a fitness advantage of these viruses on strains carrying D614G spike mutation in the hamster model. Using low-dose inocula in the hamster model, one study showed a superiority of Alpha variant when inoculated competitively and a more efficient transmission when inoculated alone[28]. This study reported that Alpha variant is slightly more excreted than G614 strain at the early stage of infection in hamsters, affecting viral transmission to contact animals. In another study that used engineered rescued viruses derived from the USA_WA1/2020 strain, the hamster model appeared useful to detect weak fitness advantages and increases in transmissibility of viruses that carry the N501Y and A570D spike mutations[9]. However, the role of other mutations located in other parts of the genome of Alpha variant [more than twenty when compared to strains isolated in January-February 2020], was not taken into account using this reverse genetics-based approach. Indeed, some studies reported that mutations located outside in the gene coding for the spike protein can also modulate fitness, transmission and virulence of SARS-CoV-2 strains[29,30].

Altogether, this suggests that the hamster model may possibly not be the best model to detect weak fitness or transmissibility differences between clinical strains of SARS-CoV-2. Other animal models such as the ferret (*Mustela putorius furo*) model that is already employed to study the pathogenicity and transmissibility of other human respiratory viruses, could be valuable tools in that case. Indeed, a recent work highlighted the importance of using multiple animal models to compare the fitness and transmission of SARS-CoV-2 strains[31]. This study failed to conclude on a clear advantage of Alpha variant over G614 strain in the hamster model, but showed a clear fitness advantage of Alpha variant over a G614 strain in ferrets and two mouse models.

## Acknowledgments

This work was supported by the European Union’s Horizon 2020 Research and Innovation Programme under grant agreements no. 653316 (European Virus Archive goes global project: http://www.european-virus-archive.com/) and also funded by REACTING/ANRS MIE. We thank Noémie Courtin for her technical support and Géraldine Piorkowski for her help on the NGS platform.

