## Supplemental figures for "Alpha variant versus D614G strain in the Syrian hamster model"

**Legends supplemental Figures and Tables:**

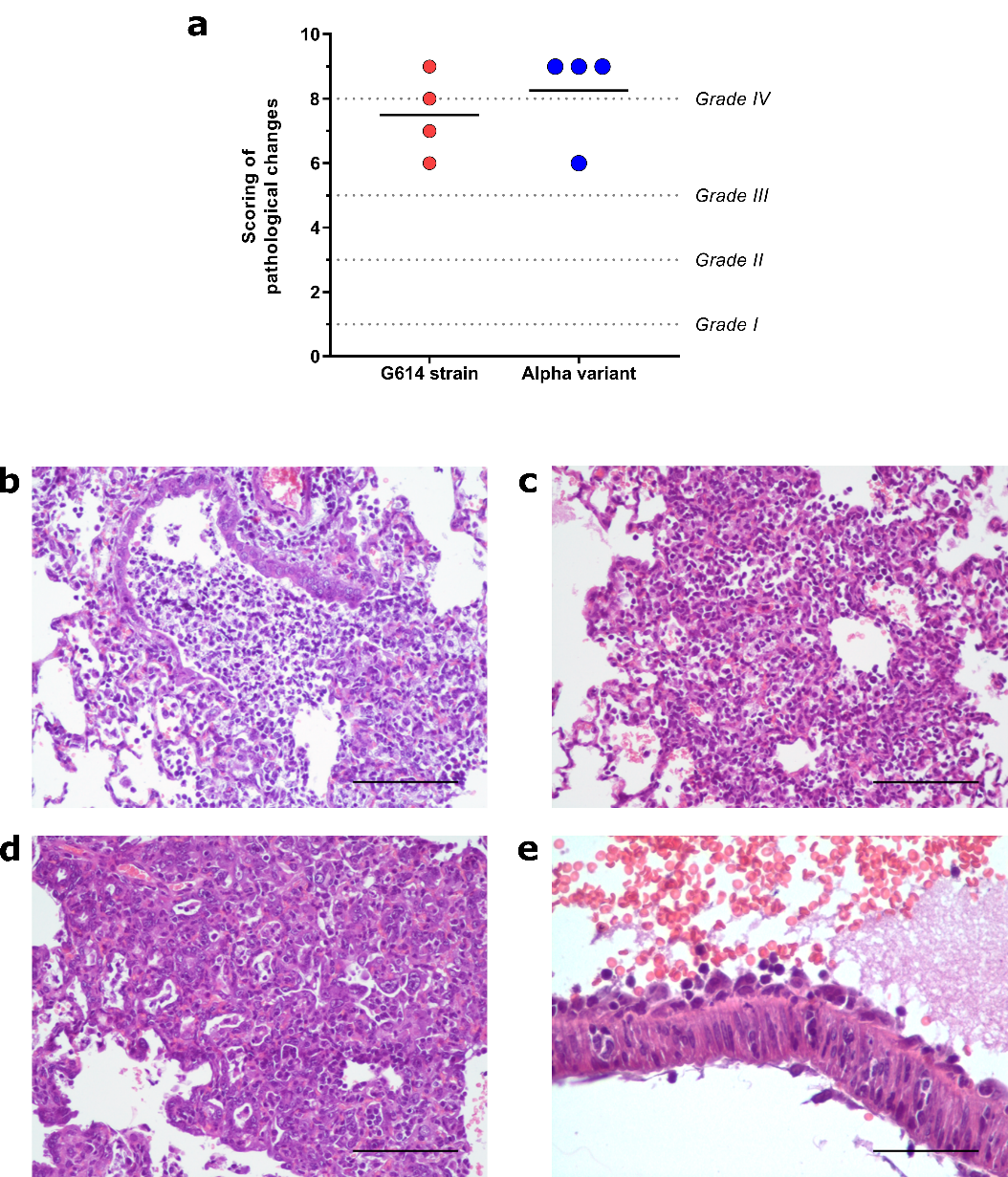

**Supplemental figure 1: Lung histopathological changes induced by Alpha variant and G614 strain.** Groups of 4 hamsters were intranasally infected with 10^4^ TCID_50_ of Alpha variant or G614 strain and sacrificed at 5 dpi. Following criteria presented in Supplemental Table 2, as previously described (*Nature Communication*. Driouich *et al*. 2021). Rows represent the mean score of each group (n=4; details in Supplemental Table 3). Difference between both groups is non-significant (Mann-Whitney test). (b-e) Representative images of lesions observed in both animal groups. (b) Representative image of bronchitis: terminal bronchiole filled with degenerated neutrophils and necrotic cellular debris (scale bar = 100µm; animal of ‘G614 strain’ group). (c) Representative images of broncho-interstitial pneumonia: area of lung consolidation with interstitial inflammation, expansion of alveolar walls by inflammatory infiltrates (macrophages, heterophils) also filling alveolar lumina (scale bar = 100µm; animal of ‘Alpha variant’ group). (d) Representative images of broncho-interstitial pneumonia: area of lung consolidation showing interstitial inflammation (macrophages, heterophils) and alveolar septa lined by low cuboïdal epithelial cell (type II pneumocytes hyperplasia) (scale bar = 100µm; animal of ‘G614 strain’ group). (e) Representative images of vasculitis, arteriolar wall infiltration by leucocytes with cellular debris, subendothelial leucocytic collection (scale bar = 50µm; animal of ‘Alpha variant’ group).

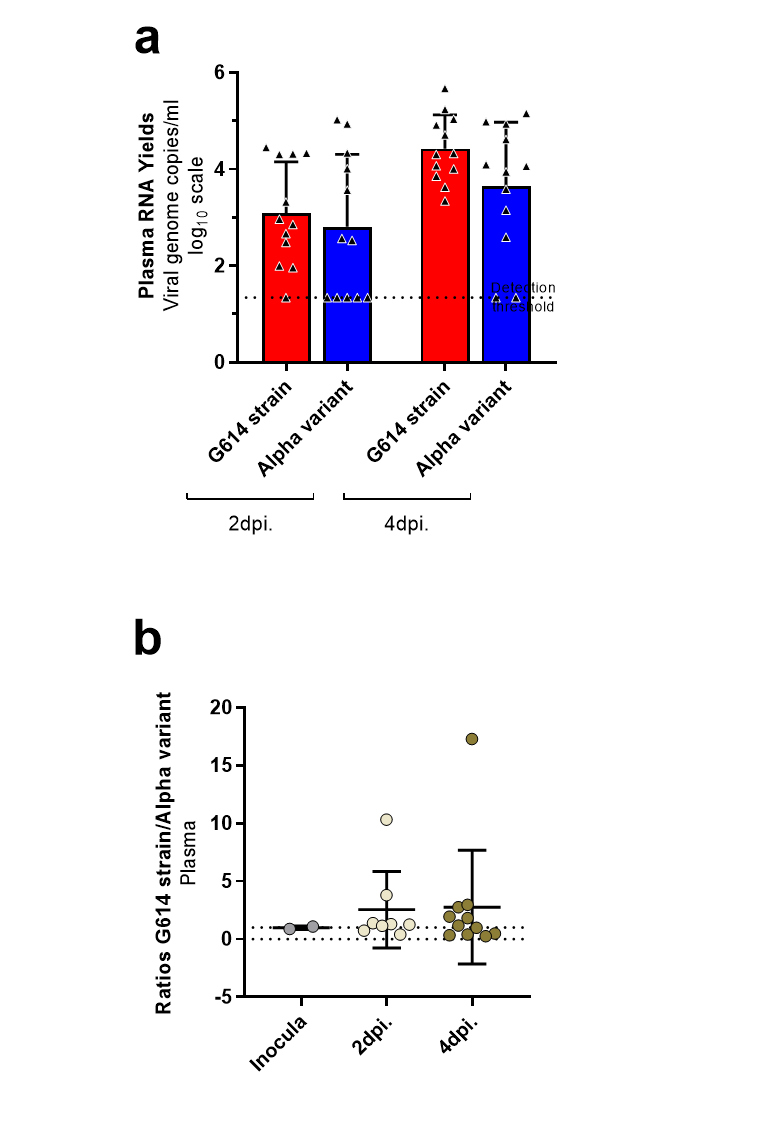

**Supplemental Figure 2: Viral RNA yields in plasmas measured using a RT-qPCR assay.** Results for (a) comparative assessment and (b) competition experiments (expressed as [G614/ Alpha] ratios). Two specific RT-qPCR assays were used to measure the quantity of each virus in plasmas.

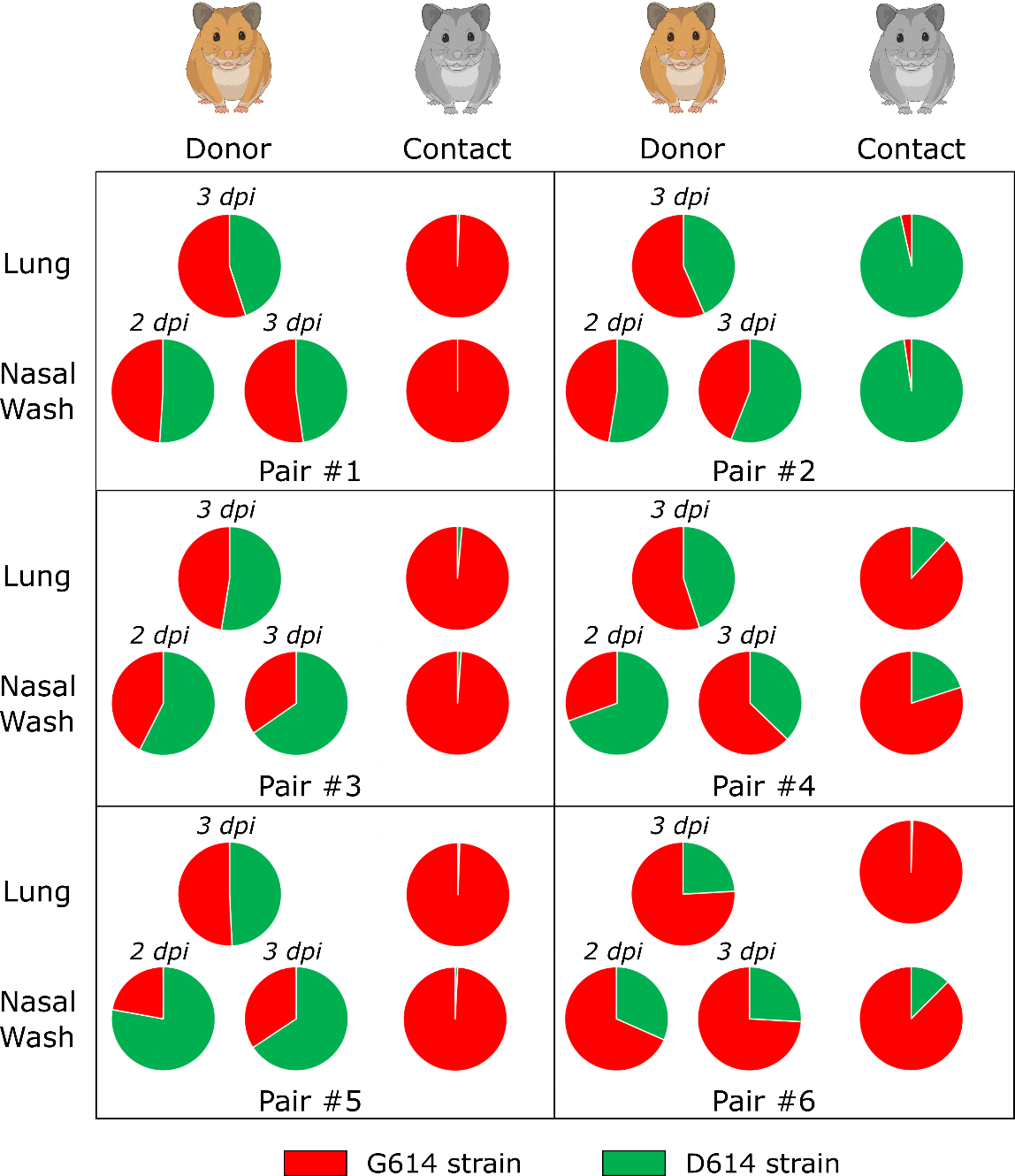

**Supplemental Figure 3:** T**ransmission experiment with G614 and D614 strains.** A group of 6 hamsters, named ‘donors’, was intranasally infected with an equal proportion of both viral strains for competition experiment (total: 20 TCID_50_). At 2 dpi, each donor was co-housed with a contact animal during a period of 6 hours and received a nasal wash. Donors and contacts were sacrificed at 3 dpi and at 3 dpc respectively. This figure is a graphical representation of the proportion of each virus found in lungs and nasal washes for each pair of animals in transmission experimentations. Results were obtained using sequencing method described in the ‘Molecular biology’ part of the ‘Materiel and Methods’ section.

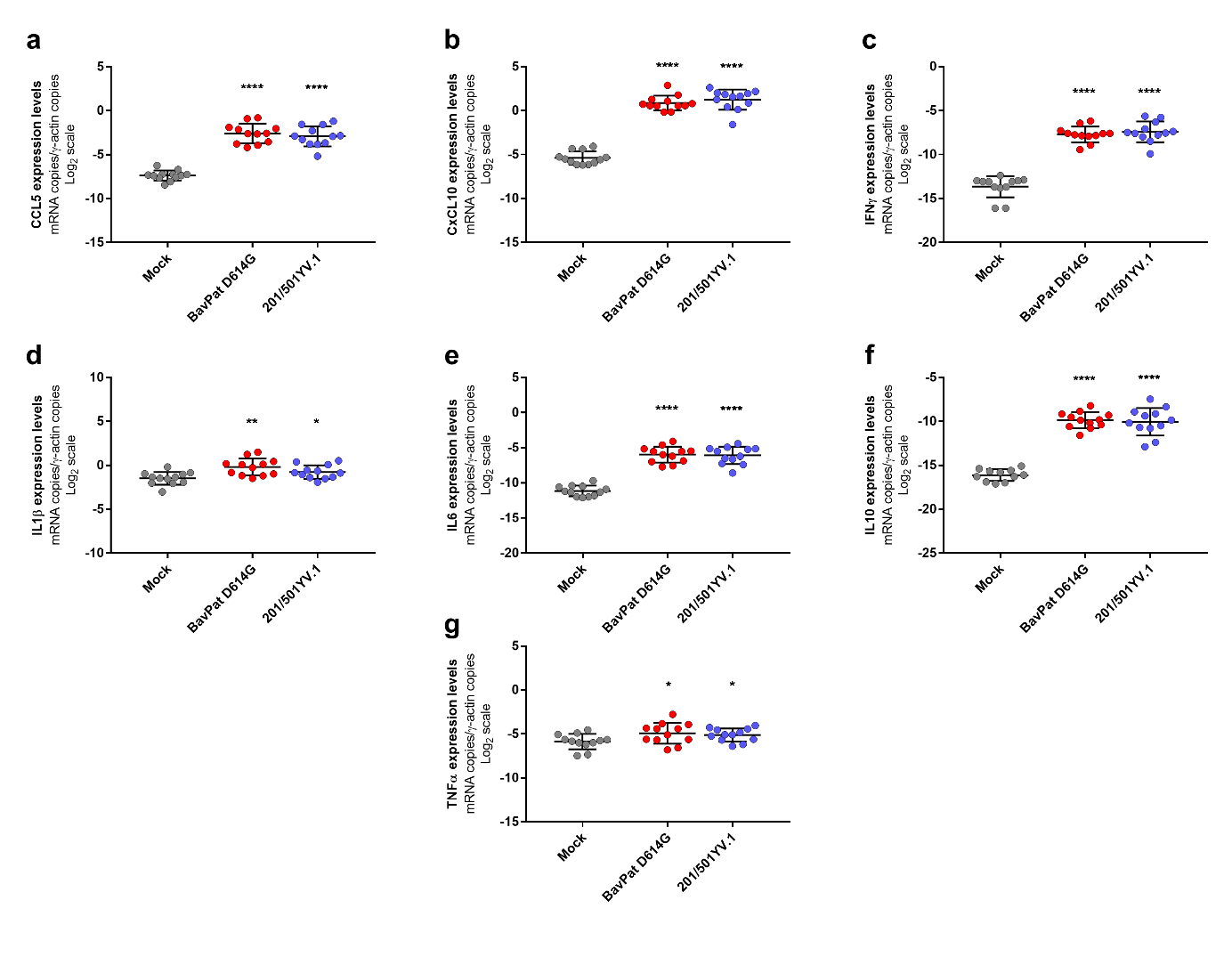

**Supplemental Figure 4:** **Expression level of cytokines genes in the lungs at 4 dpi after infection with G614 strain or Alpha variant.** A panel of seven cytokines is presented: Ccl5 (a), Cxcl-10 (b), Interferon-γ (c), IL-1β (d), IL-6 (e), IL-10 (f) and TNF-α (g). Gene expression was quantified using RT-qPCR assays expressed as mRNA copies/γ-actin copies. ***, ** and * symbols indicate that the mean expression for the G614 or for the Alpha group are significantly higher than those of the mock-infected group with a p-value ranging between 0.0001-0.001, 0.001–0.01, and 0.01–0.05, respectively (Mann-Whitney and unpaired t tests).
